# Benzo[a]pyrene induces keratinocyte senescence and p21-dependent differentiation

**DOI:** 10.64898/2026.05.08.723713

**Authors:** Cheng Lui Daniel Law, Li Fang Melissa Tang, Maurice A.M. van Steensel

## Abstract

1. In this study, we demonstrate that Benzo[a]pyrene (B[a]P) induces keratinocyte senescence and p21^Cip1^-dependent keratinocyte differentiation. Atmospheric and environmental pollution are known to induce senescence and promote terminal differentiation in human primary keratinocytes, thus driving skin aging. However, much is still unknown about the underlying molecular mechanisms. We observed that B[a]P, a common atmospheric pollutant, induced senescence in primary keratinocytes in both two-dimensional and three-dimensional (reconstructed human epidermis) culture. This was accompanied by signs of DNA damage in B[a]P-treated cells. B[a]P-treated cells also underwent accelerated late-stage terminal differentiation, indicated by increased *IVL* and *FLG* expression from 48 to 96 hours post-exposure. While pharmacological and genetic attenuation of p21^Cip1^ did not rescue cellular senescence, it prevented the expression of *IVL* and *FLG*, suggesting that the late-stage terminal differentiation induced by B[a]P exposure was p21-dependent. Our data thus suggest a key role for the p21^Cip1^ in the keratinocyte response to pollution-induced damage, where p21^Cip1^ induces terminal differentiation to maintain skin barrier homeostasis.

## INTRODUCTION

Skin aging can be divided into two distinct processes: intrinsic and extrinsic aging. Intrinsic/chronological skin aging is driven by the accumulation of senescent cells (Ho and Dreesen, 2021). Extrinsic skin aging arises from chronic damage due to environmental factors (Zhang and Duan, 2018), and can further accelerate the formation of damaged and senescent cells. These include, but are not limited to, ultraviolet irradiation, air pollution and tobacco smoke (Krutmann et al., 2017), culminating in the premature stress-induced senescence of skin cells and loss of skin barrier homeostasis. Indeed, cellular senescence is a major hallmark of aging, and the role of extrinsic factors in exacerbating cellular senescence is well established (Biniek et al., 2012, Kim et al., 2021, Ryu et al., 2019a, Tan et al., 2022).

Senescence is characterised by a state of permanent cell cycle arrest, which is induced by activation of the p53/p21^Cip1^ pathways and p16^INK4a^/Retinoblastoma pathways (Alcorta et al., 1996, Beausejour et al., 2003, Stein et al., 1999, Sturmlechner et al., 2021). Extrinsic stressors can induce oxidative stress and DNA damage in skin cells. Subsequently, the p53-p21^Cip1^ pathway is activated, resulting in cell cycle arrest and senescence (He et al., 2005). For example, p21^Cip1^ upregulation is associated with UVB-induced senescence in melanocytes and murine keratinocytes (Liu and Pelling, 1995, Martic et al., 2020), and the p53-p21^Cip1^ pathway is also activated upon H_2_O_2_-induced keratinocyte senescence (Ido et al., 2012).

Air pollution, specifically particulate matter (PM), has been reported to cause accelerated skin aging (Huls et al., 2016, Vierkotter et al., 2010). Importantly, polycyclic aromatic hydrocarbons (PAHs) are a significant toxic component that is typically found adsorbed to PM and have been demonstrated to be highly genotoxic to lung and foreskin fibroblasts (Allmann et al., 2020, Oh et al., 2011, Xu et al., 2024). Previously, PM was demonstrated to induce keratinocyte senescence through oxidative stress-induced upregulation of p16^INK4A^ expression (Ryu et al. 2019). However, the mechanistic basis of pollution-induced skin aging is still unclear.

The PAH Benzo[a]Pyrene (B[a]P) is one of the main components of air and environmental pollution (Bukowska et al., 2022, Hattemer-Frey and Travis, 1991) and is a widely used tool compound for air or environmental pollution research (Allmann et al., 2020, Billiard et al., 2006, Tsuji et al., 2011). B[a]P is ubiquitous in the environment and is classified as a Group 1 carcinogen by the International Agency for Research on Cancer (Baan et al., 2009). It is metabolised to the genotoxic intermediate benzo[a]pyrene diolepoxide (BPDE) in the cytosol by CYP1A1 (Yang et al., 1977), where it forms a bulky BPDE-DNA adduct that can induce double-stranded (ds)DNA breaks (Allmann et al., 2020). Additionally, B[a]P is an extremely potent AhR ligand (Giani Tagliabue et al., 2019, Hidaka et al., 2017, Zhou et al., 2017). AhR activation has been demonstrated to induce the release of inflammatory cytokines from keratinocytes (Afaq et al., 2009, Qiao et al., 2017, Ryu et al., 2019b). Additionally, AhR-induced ROS accumulation can indirectly induce human primary keratinocyte (HPK) senescence (Ryu et al., 2019a). However, to our knowledge, a role for B[a]P in skin aging has not been investigated yet.

In this study, we used *in vitro* keratinocyte culture and reconstructed human epidermis models to demonstrate the ability of B[a]P, a well-established air pollutant, to drive HPK senescence and differentiation. We show that p21^Cip1^ is crucial for the regulation of B[a]P-induced HPK senescence and differentiation and demonstrate how a partial AhR knockdown results in the rescue of B[a]P-induced HPK senescence. Finally, we used pharmacological inhibition and genetic knockdowns to further investigate the importance of p21^Cip1^ for late-stage keratinocyte differentiation, where we demonstrated that the inhibition of p21^Cip1^ by both pharmacological and genetic means resulted in a delay of B[a]P-induced late-stage keratinocyte differentiation.

## RESULTS

### Benzo[a]pyrene induces cellular senescence in keratinocytes in both 2D keratinocyte and 3D reconstituted human epidermis models

Previously, diesel particulate matter (DPM) was shown to induce senescence in HPKs through the upregulation of p16^INK4A^ (Ryu et al., 2019a). We expanded on this model of keratinocyte senescence by looking at various established biomarkers of senescence. Both H_2_O_2_ and DPM treatment resulted in a higher proportion of larger and flatter cells that stained positively for SA-β-Gal activity (Figure S1a). Lamin B1 (LMNB1) expression was decreased at both the protein and transcript level (Figure S1b, S1d) while DPM-treated cells demonstrated an overall increase in *CDKN1A* (p21^CIP1^) gene expression, albeit significant only at 25μg/ml DPM (Figure S1b). Since air and environmental pollution were reported to activate the AhR signalling pathway (Afaq et al., 2009, Krutmann et al., 2014, Qiao et al., 2017), we decided to interrogate expression of *AhR* and an established AhR target gene, *CYP1A1* (Ma, 2001). Indeed, *CYP1A1* gene expression was significantly increased, indicating the activation of the AhR pathway (Figure S1c). In addition, we observed fewer keratinocytes staining positively for Ki67, suggesting loss of proliferative capacity (Figure S1d). Collectively, these observations indicated that DPM induced senescence in keratinocytes in a dose-dependent fashion.

DPM has been used in the past to study the effects of particulate matter pollution (Lee et al., 2021, O’Driscoll et al., 2018, Ryu et al., 2019a). However, DPM is of limited utility for mechanistic studies because of its heterogenous nature, consisting of a mixture of polycyclic aromatic hydrocarbons (PAHs) such as Benzo[a]pyrene (B[a]P), nitrogen containing-PAHs, heavy metals and soot. There are also strong batch variations, further confounded by the geographic location where the DPM was collected, which influences its composition (Liu et al., 2024). Hence, with DPM it would be difficult to tease out any specific mechanisms driving the keratinocyte response to pollution. Thus, we decided to use B[a]P to model pollution-induced keratinocyte senescence.

With 200μM H_2_O_2_-treated HPKs as a senescent control, we found that exposure to 2μM B[a]P resulted in a larger proportion of SA-β-Gal positive keratinocytes (Figure 1a) compared to non-treated controls. In contrast, 20nM 2,3,7,8-Tetrachlorodibenzodioxin (TCDD), a highly potent and selective AhR agonist (Giani Tagliabue et al., 2019, Ma and Baldwin, 2000), did not induce senescence in keratinocytes. Furthermore, gene and protein expression of LMNB1in B[a]P-treated cells were almost completely lost in both H_2_O_2_ and B[a]P treated keratinocytes but were sustained in TCDD-treated keratinocytes (Figure 1b-d). This was accompanied by a sharp reduction in the number of Ki67-positive cells in only H_2_O_2_ and B[a]P-treated keratinocytes (Figure 1d). *CDKN1A* (p21^Cip1^) gene expression did not significantly change after 5 days of B[a]P exposure (Figure 1c). Finally, we validated B[a]P-induced senescence in Reconstructed Human Epidermis (RHE), where 2μM B[a]P was topically applied every 2 days up till day 7. While H&E staining did not reveal any significant morphological defects (Supplementary Figure S2), the total expression of LMNB1 in the RHE was reduced drastically post-treatment (Figure 1e). Taken together, these observations led us to conclude that B[a]P, but not TCDD, induced senescence in primary keratinocytes and that AhR signalling was not involved in B[a]P-induced keratinocyte senescence.

**Figure 1:**
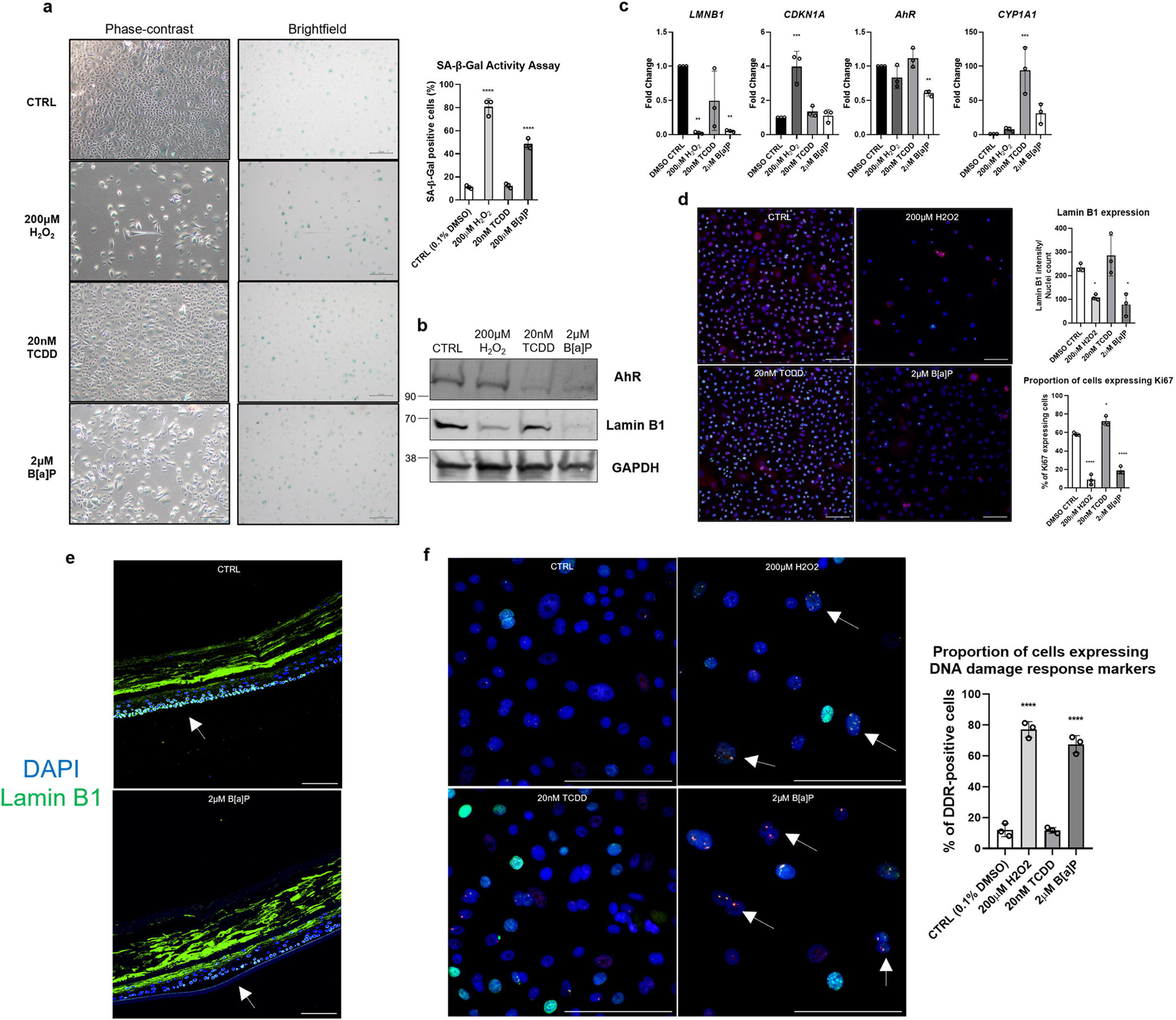
Benzo[a]pyrene induced cellular senescence in primary keratinocytes and Reconstructed Human Epidermis. (a) SA-β-Gal staining and imaging of HPKs (2D cultures) performed after treating HPKs with 200μM H_2_O_2_ for 1h and allowing 5 days for recovery, and 20nM TCDD and 2μM B[a]P for 5 days. Bar = 200μM. (b) Western Blot analyses after treating HPKs as in a. (c) RT-qPCR after treating HPKs as in a. (d) Immunofluorescence analyses (Lamin B1, Red; Ki67, Green; DAPI, Blue) was performed after treating HPKs as in a. Bar = 100μm (e) Immunofluorescence analyses (Lamin B1, Green; DAPI, Blue) of Reconstructed Human Epidermis topically treated with 2μM B[a]P after 7 days. (f) Immunofluorescence analyses (53BP1, Red; γH2AX, Green; DAPI, Blue) was performed after treating HPKs with 200μM H_2_O_2_ for 1h and allowed 5 days for recovery, and 20nM TCDD and 2μM B[a]P for 5 days. White arrows indicate DNA damage foci. Bar = 100μm. All treatments were compared to DMSO control for statistical testing. Statistical analyses: One-way ANOVA, n = 3, *P*-values < 0.05 indicate statistical significance (∗P < 0.05; ∗∗P < 0.01; ∗∗∗P < 0.001; ∗∗∗∗P < 0.0001). SA-β-Gal (Senescence-Associated Beta Galactosidase); H_2_O_2_ (Hydrogen peroxide); TCDD (2,3,7,8-Tetrachlorodibenzodioxin); B[a]P (Benzo[a]pyrene); *LMNB1* (Lamin B1); *CDKN1A* (p21^Cip1^).

To probe the mechanism by which B[a]P induces senescence, we analyzed AhR pathway activation. Both B[a]P and TCDD activated the AhR pathway, based on their induction of *CYP1A1* gene expression by about 30- and 100-fold, respectively. In addition, AhR was completely degraded in AhR agonist-treated keratinocytes, further supporting AhR pathway activation (Figure 1b) as described previously (Ma and Baldwin, 2000). Interestingly, only B[a]P exposure resulted in the significant loss of *AhR* gene expression by about two-fold (Figure 1c). Upon H_2_O_2_ and B[a]P exposure, we observed sustained accumulation of γH2AX/53BP1 DNA damage foci in the nuclei of keratinocytes, indicating double-stranded DNA damage (Lassmann et al., 2010, Pessina et al., 2019, Popp et al., 2017, Williamson et al., 2020).. Notably, TCDD did not induce any accumulation of γH2AX/53BP1 foci (Figure 1f). In summary, while both B[a]P and TCDD activated the AhR pathway, only B[a]P induced DNA damage and senescence.

### Benzo[a]pyrene induced primary keratinocyte differentiation

To further characterize the mechanism(s) underlying B[a]P-induced senescence, we compared the transcriptomes of untreated controls with those of B[a]P-exposed keratinocytes. We identified a total of 1432 significantly differentially expressed genes (DEGs) (defined by adjusted *P* value < 0.05 and absolute fold-change ≥ 2), with 849 genes upregulated and 583 genes downregulated (Supplementary Figure S3). We used Gene Ontology (GO) enrichment analysis to obtain an overview of pathways that might be differentially regulated. Interestingly, it revealed that multiple GO terms associated with terminal keratinocyte differentiation were amongst the most significantly upregulated. This strongly suggested that

B[a]P treatment induced late-stage terminal keratinocyte differentiation, (Figure 2a). Subsequently, Gene Set Enrichment Analysis (GSEA) was performed on GO terms associated with keratinocyte differentiation to characterize the keratinocyte differentiation phenotype (Figure 3b). Remarkably, all keratinocyte differentiation gene sets presented as significantly enriched (cutoff FDR < 0.25) in B[a]P-treated keratinocytes.

**Figure 2:**
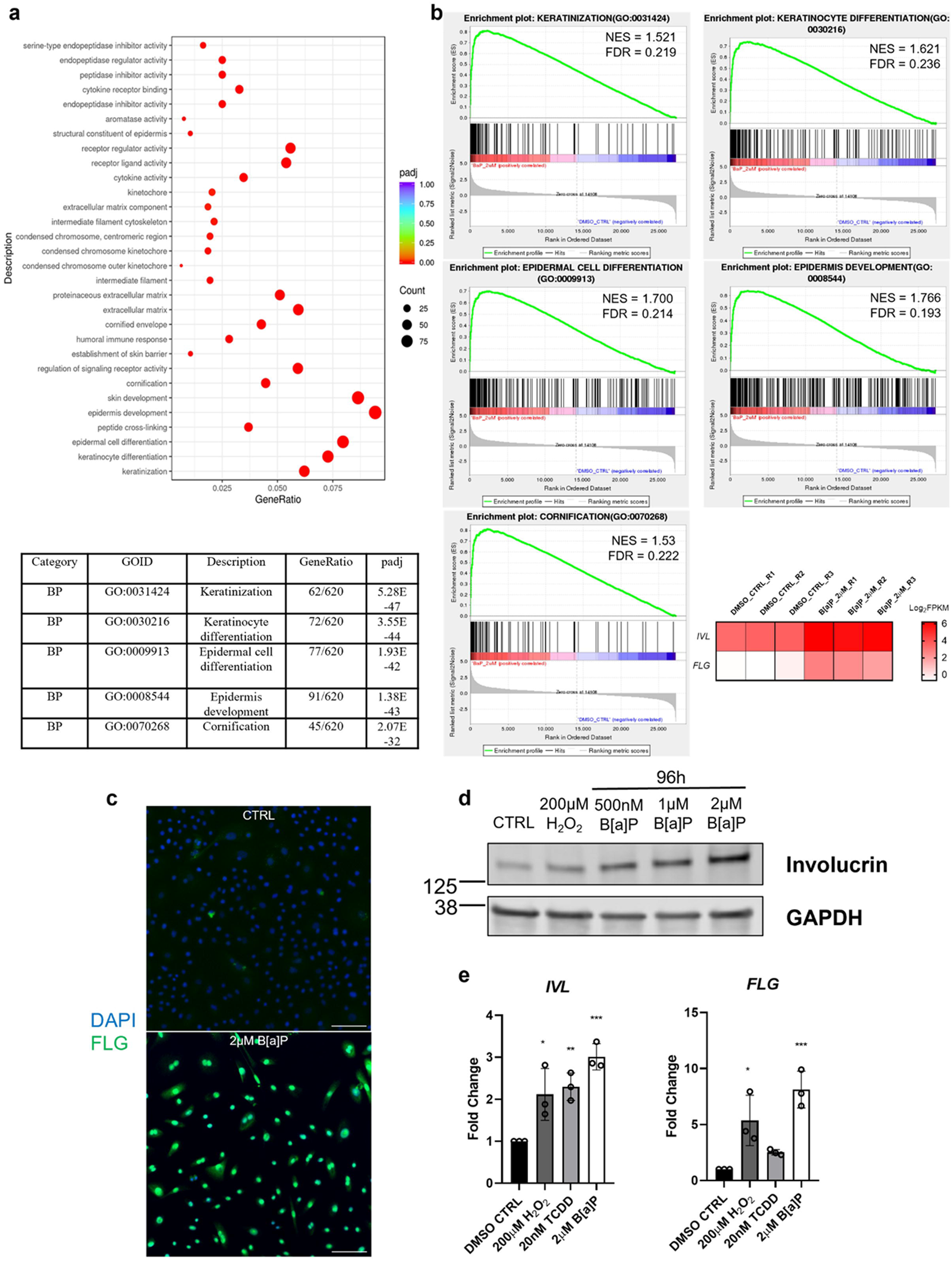
Benzo[a]pyrene induced primary keratinocyte differentiation. (a) DMSO CTRL vs 2μM B[a]P HPK treatment for 4 days. GO Pathway Enrichment bubble plot: The abscissa in the graph is the ratio of the differential gene number to the total number of differential genes on the GO Term, and the ordinate is GO Term. 30 most significant terms were listed, and adjusted *P* values for GO terms associated to keratinocyte differentiation listed in the table below. (b) GSEA individual gene set enrichment plots for GO terms that are related to keratinocyte differentiation are described here, along with heatmap depicting *IVL* and *FLG* expression. (c) Immunofluorescence analyses (Filaggrin, Green; DAPI, Blue) was performed after treating HPKs with 2μM B[a]P for 4 days. Bar = 100μm (d) Western Blot analyses was performed after treating HPKs with 200μM H_2_O_2_ for 1h and allowed 4 days for recovery, and 500nM-2μM B[a]P for 4 days. (e) RT-qPCR analyses performed after treating HPKs with 200μM H_2_O_2_ for 1h and allowed 5 days for recovery, and 20nM TCDD and 2μM B[a]P for 5 days. Statistical analyses: One-way ANOVA, n = 3, *P*-values < 0.05 indicate statistical significance (∗P < 0.05; ∗∗P < 0.01; ∗∗∗P < 0.001; ∗∗∗∗P < 0.0001). H_2_O_2_ (Hydrogen peroxide); TCDD (2,3,7,8-Tetrachlorodibenzodioxin); B[a]P (Benzo[a]pyrene); *IVL* (Involucrin); *FLG* (Filaggrin).

**Figure 3:**
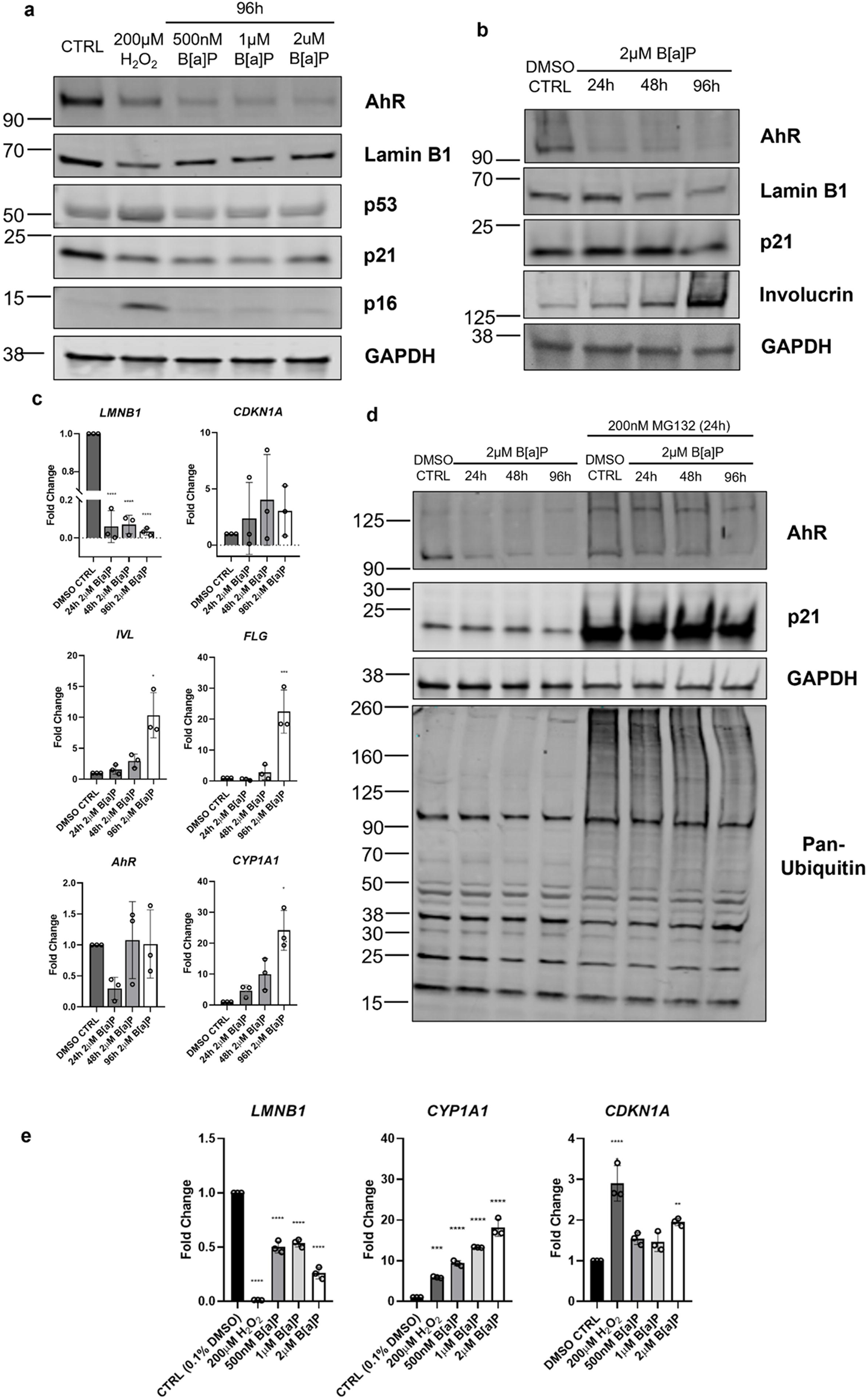
p21^Cip1^ expression is reduced in B[a]P-induced keratinocyte senescence and differentiation. (a) Western Blot analyses was performed after treating HPKs c, and 500nM-2μM B[a]P for 4 days. (b) Western Blot analyses was performed after treating HPKs with 2μM B[a]P for 1(24h), 2(48h) and 4(96h) days. (c) RT-qPCR analyses was performed after treating HPKs as described in b. (d) Western Blot analyses was performed after treating HPKs with 2μM B[a]P for 1(24h), 2(48h) and 4(96h) days with or without 200nM MG132 on the final 24 hours of treatment. (e) RT-qPCR analyses after treating HPKs with 200μM H_2_O_2_ for 1 hour and allowed 4 days to recover and 500nM/1μM/2μM B[a]P for 4 days. Statistical analyses: One-way ANOVA, n = 3, *P*-values < 0.05 indicate statistical significance (∗P < 0.05; ∗∗P < 0.01; ∗∗∗P < 0.001; ∗∗∗∗P < 0.0001). H_2_O_2_ (Hydrogen peroxide); B[a]P (Benzo[a]pyrene); *LMNB1* (Lamin B1); *CDKN1A* (p21^Cip1^); *IVL* (Involucrin); *FLG* (Filaggrin).

As GO terms associated with keratinocyte differentiation, *IVL* and *FLG* were included within all gene sets investigated by GSEA (Dale et al., 1985, Eckert and Rorke, 1989). As expected, both *IVL* and *FLG* gene expression were enhanced sharply in B[a]P-treated keratinocytes (Figure 2b). Orthogonal validation was then carried out using multiple methods to validate elevated mRNA and protein expression of both IVL and FLG (Fig. 2c and 2d). Curiously, TCDD treatment induced a significant increase in *IVL* transcripts compared to the control (Fig. 2e). Hence, AhR signalling might be partially responsible for the upregulation of late-stage terminal differentiation genes. Taken together, the evidence suggested that in addition to senescence, B[a]P might also promote the expression of cornified envelope proteins such as IVL and FLG in keratinocytes, suggesting a potential role in inducing keratinocyte terminal differentiation.

### p21^Cip1^ expression is reduced in B[a]P-induced keratinocyte senescence and differentiation

The p53/p21^Cip1^pathway and p16^INK4a^/pRB pathway are the major drivers of the cell cycle arrest observed in senescence (Campisi, 2013). Senescence of skin cells such as keratinocytes and fibroblasts cultured *in vitro*, in reconstructed skin and skin explants has been identified through observing the elevated expression of p53, p21^Cip1^ and p16^INK4a^ as a biomarker (Fitsiou et al., 2021, Tan et al., 2022, Wang et al., 2022). Hence, to study if the same pathways were involved in B[a]P-induced senescence, we looked at p53, p21^Cip1^ and p16^INK4a^ protein expression. Both H_2_O_2_ and 4 days of B[a]P exposure induced senescence as seen by the reduction of lamin B1 at both protein and mRNA levels, while B[a]P strongly induced AhR activation (Figure 3a, 3c, 3e). H_2_O_2_-exposure induced sustained p53 and p16^INK4a^ accumulation, suggesting involvement of p53/p21^Cip^ and p16^INK4a^ pathways in H_2_O_2_-induced senescence. Surprisingly, both p53 and p16^INK4a^ expression were unchanged compared to the

DMSO control after 4 days of B[a]P exposure, implying that sustained p53 and p16^INK4a^ expression were not necessary for the maintenance of B[a]P-induced senescence (Figure 3a). Conversely, we observed a stark reduction of p21^Cip1^ expression as keratinocytes became senescent (Figure 3a). Taken together, our observations suggest a potential association between p21^Cip1^ and B[a]P-induced HPK senescence.

We proceeded to perform a time-course experiment to further elucidate the kinetics of p21^Cip1^ expression in B[a]P-treated keratinocytes. We found that over the course of 4 days, LMNB1 and IVL expression gradually decreased and increased, respectively, demonstrating that B[a]P-treated HPKs were indeed entering senescence and terminal differentiation. Notably, p21^Cip1^ expression levels were slightly increased for the first two days and only dipped after the 4^th^ day (Figure 3b). This was accompanied by a strong time-dependent increase in gene expression of *FLG* and *IVL* (Figure 3c). These observations suggested to us that p21^Cip1^ may play a role in the regulation of terminal differentiation through the modulation of cornified envelope genes such as *FLG* and *IVL*. Hence, we hypothesized that p21^Cip1^is necessary for the induction of the terminal differentiation programme.

We observed that while p21^Cip1^ protein expression was reduced after 4 days of B[a]P exposure, p21^Cip1^ gene expression was upregulated significantly, suggesting p21^Cip1^ expression was regulated via post-translational mechanism (Figure 3e). Past reports have demonstrated 20S proteasomal-mediated degradation of p21^Cip1^ (Bloom et al., 2003, Lu and Hunter, 2010). Hence, we decided to test whether the observed degradation of p21^Cip1^ required the ubiquitin-proteasomal pathway. Indeed, inhibition of the 20S proteasome with MG132 rescued p21^Cip1^ protein expression (Figure 3d). Our results suggest that B[a]P exposure resulted in the ubiquitin-proteasomal degradation of p21^Cip1^ after 4 days, coinciding with the strong induction of IVL expression. Thus, there may be an association between p21^Cip1^ and terminal differentiation in HPKs.

### Pharmacological attenuation and genetic knockdown of p21^Cip1^ both delay B[a]P-induced late-stage keratinocyte differentiation

To elucidate p21^Cip1^’s role in terminal differentiation, we used UC2288, a small molecule that inhibited p21^Cip1^ at both the transcript and protein levels (Wettersten et al., 2013). Interestingly, the partial attenuation of p21^Cip1^ by UC2288 inhibited IVL expression, but did not rescue the loss of LMNB1, indicating that the loss of p21^Cip1^ may have inhibited or slowed late-stage terminal differentiation of keratinocytes, but did not affect senescence (Figure 4a). While keratinocyte differentiation and senescence are generally observed to occur concurrently in response to external stress such as UVB irradiation (Tan et al., 2019, Tan et al., 2022), our observations suggest that these two processes are distinct and regulated by different pathways.

**Figure 4:**
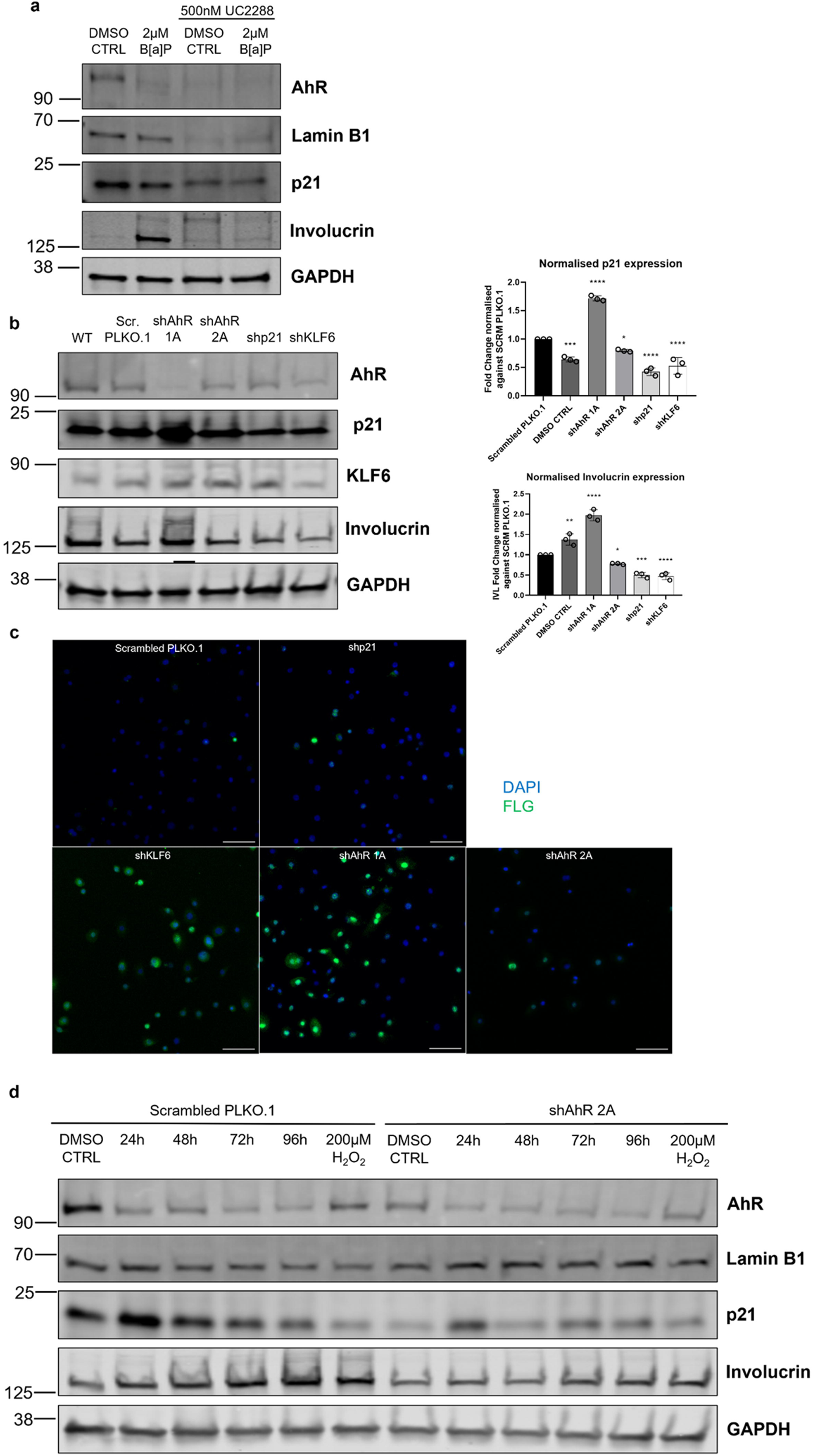
Pharmacological attenuation and genetic knockdown of p21^Cip1^ results in the delay of B[a]P-induced late-stage keratinocyte differentiation. (a) Western Blot analyses was performed after treating HPKs with 2μM B[a]P for 4 days, with or without 500nM UC2288. (b) Western Blot analyses of near complete shRNA-mediated knockdown of AhR (shAhR 1A), Partial shRNA-mediated knockdown of AhR (shAhR 2A), Partial shRNA-mediated knockdown of p21^Cip1^ (shp21), Partial shRNA-mediated knockdown of KLF6 (pKLF6) was carried out 4 days post-lentiviral transduction. Quantification of relative expression against the control (Scrambled PLKO.1) for p21^Cip1^ and involucrin was performed. (c) Immunofluorescence analyses (Filaggrin, Green; DAPI, Blue) was carried out as described in b. Bar = 100μm. (d) Western Blot analyses of shAhR 2A vs scrambled PLKO.1 was performed after treating HPKs with 200μM H_2_O_2_ for 1h and allowed 4 days for recovery, and 2μM B[a]P for 1-4 days. H_2_O_2_ (Hydrogen peroxide); B[a]P (Benzo[a]pyrene); *FLG* (Filaggrin). Statistical analyses: One-way ANOVA, n = 3, *P*-values < 0.05 indicate statistical significance (∗P < 0.05; ∗∗P < 0.01; ∗∗∗P < 0.001; ∗∗∗∗P < 0.0001).

Importantly, p21^Cip1^ expression is known to be regulated by the non-canonical AhR-KLF6 transcription activating complex, which binds to non-consensus xenobiotic-response elements (NC-XREs). (Wilson et al., 2013). Thus, to further investigate if p21^Cip1^ is a key driver of late-stage keratinocyte differentiation, we performed shRNA-mediated knockdown of *AhR*, *CDKN1A*, and *KLF6*. Indeed, transduction of shAhR 2A (less efficient KD of AhR), as well as shp21 and shKLF6 (partial KD of *CDKN1A* and *KLF6* respectively) resulted in varying levels of p21^Cip1^ reduction. Paradoxically, near complete KD of *AhR* (with shAhR 1A) resulted in drastic accumulation of p21^Cip1^ (Figure 4b). More importantly, IVL expression correlated with the level of p21^Cip1^ expression. When p21^Cip1^ was highly expressed in shAhR 1A-transduced HPKs, a corresponding increase of IVL expression was observed. However, when p21^Cip1^ expression was reduced in shAhR 2A, shp21 and shKLF6-transduced HPKs to varying extents, we observed a corresponding reduction of IVL expression (Figure 4b). Thus, although we do not have direct mechanistic evidence, our observations suggest that p21^Cip1^ may have a role in inducing IVL expression. To further validate this observation, we investigated FLG expression in the genetically modified keratinocytes. We observed that shp21 and shAhR 2A-transduced HPKs had similar FLG expression to the scrambled control. However, shAhR 1A and shKLF6-transduced HPKs spontaneously differentiated in culture and demonstrated a robust upregulation of FLG expression (Figure 4c). These observations mostly supported our observations of p21^Cip1^ being associated with the modulation of HPK terminal differentiation. We hypothesize that, while p21^Cip1^ may drive keratinocyte differentiation and IVL expression, FLG might be also regulated by other pathways.

As the other shRNA-transduced cells did not proliferate, the only stable shRNA-transduced cell line we could establish was shAhR 2A. To further characterise the role of AhR in B[a]P - induced senescence and differentiation in HPKs, we performed a B[a]P exposure time-course experiment for 1-4 days to compare p21^Cip1^ kinetics between shAhR 2A-transduced HPKs and the scrambled control. Interestingly, in shAhR 2A-transduced keratinocytes, p21^Cip1^ expression was also severely reduced. Similar to the scrambled control, p21^Cip1^ was sharply upregulated after 1 day of B[a]P exposure in shAhR 2A-transduced keratinocytes. This was followed by a loss of p21^Cip1^ until after the 4^th^ day. We also observed that shAhR 2A-transduced keratinocytes were resistant to B[a]P-induced senescence as indicated by the rescue of LMNB1 expression and reduction of SA-β-Gal positive cells (Figure 4d and Supplementary Figure S4). In addition, delayed differentiation was observed, as seen from the relative decrease of IVL expression even after the 4^th^ day of B[a]P exposure (Figure 4d).

## DISCUSSION

In this study, we report that the ubiquitous environmental pollutant Benzo[a]pyrene (B[a]P), a potent carcinogen, induced DNA damage, senescence and accelerated late-stage terminal differentiation in human primary keratinocytes. We also observed that HPK terminal differentiation and senescence are two distinct processes. Keratinocytes exposed to 2 µM B[a]P accumulated p21^Cip1^ for the first 24 to 48 hours, most likely to induce cell cycle arrest in response to the extensive double-stranded DNA damage induced by B[a]P. Following this, keratinocytes entered senescence, which coincided with a commitment to terminal differentiation as observed from the upregulation of IVL expression. Subsequently, only after the differentiation phenotype was strongly induced and sustained, p21^Cip1^ was rapidly degraded by the ubiquitin-proteasomal system. Upon inhibiting p21^Cip1^ expression using both pharmacological and genetic approaches, we observed that involucrin expression was greatly decreased even after 96 hours of B[a]P exposure, demonstrating that p21^Cip1^ regulates involucrin expression and, thus, at least some aspects of keratinocyte terminal differentiation. Furthermore, we observed that the accumulation of p21^Cip1^ in the first 24 hours was associated with a sharp increase of involucrin and filaggrin expression at 96 hours, suggesting that p21^Cip1^ may have a role in keratinocyte terminal differentiation.

Taken together, our data suggests that p21^Cip1^ is an important decision-making node for pollution-induced HPK terminal differentiation. We hypothesize that in keratinocytes, the response to pollution-induced skin damage is globally integrated into two major outcomes: senescence and differentiation. These two outcomes likely aid to prevent cellular transformation occurring in a damaged keratinocyte, and to reinforce the skin barrier. Additionally, these integrated responses most likely serve to maintain or enhance the skin barrier in the face of external damage. Indeed, in chronologically aged, pollution-damaged and photoaged skin, the skin barrier is typically compromised (Biniek et al., 2012, Kim et al., 2021, Lee et al., 2016, Wang et al., 2020). This may be because senescent basal keratinocytes that were already damaged beyond repair might be driven into terminal differentiation to bolster the epidermal skin barrier to maintain skin barrier homeostasis. In summary, pollution-induced skin aging may result in the dysregulation of senescence and differentiation, possibly culminating in stem cell niche depletion.

Another interesting observation was that AhR expression levels seemed to influence the sensitivity of keratinocytes to B[a]P-induced senescence and differentiation. When AhR was partially knocked down (shAhR 2A-transduced HPKs), we observed a corresponding loss of p21^Cip1^ expression and a delay in B[a]P-induced keratinocyte senescence and differentiation. This could be due to the reduction of genotoxicity that arises from the conversion of B[a]P to BPDE by CYP1A1. We also observed less p21^Cip1^ accumulation, which led to reduced senescence and HPK differentiation. Indeed, we observed rescued levels of Lamin B1 and reduced levels of IVL in partial AhR knockdown keratinocytes after B[a]P treatment compared to the scrambled control (Figure 4d). Conversely, when AhR was nearly completely knocked down (shAhR 1A), p21^Cip1^ accumulated in keratinocytes and sharply upregulated the expression of involucrin and filaggrin even without B[a]P exposure (Figure 4b). In addition, these keratinocytes spontaneously went into cell cycle arrest and initiated terminal differentiation (Figure 4b and c). Thus, we hypothesize that AhR activation could be essential for the maintenance of cellular proliferation, and that AhR’s near-complete reduction drives keratinocyte senescence and terminal differentiation. Indeed, AhR pathway activation induced by coal tar application has been shown to induce skin barrier repair in atopic dermatitis (AD) patients (Han et al., 2016, van den Bogaard et al., 2013, Yin et al., 2016). Thus, regulation of AhR signalling likely takes place in a Goldilocks zone: strong AhR activation results in accelerated keratinocyte senescence and terminal differentiation, while the near complete knockdown of AhR in keratinocytes leads to cessation of proliferation and rapid terminal differentiation. Similar to how coal tar is used to drive keratinocyte differentiation and barrier formation to combat skin barrier dysfunction in AD, we propose that active compounds that can bring pollution-induced AhR signalling back down to normal physiological levels might ameliorate pollution-induced skin senescence and differentiation.

In conclusion, our data strongly implies that B[a]P-induced DNA damage induces senescence and subsequent differentiation in HPK. Thus, when keratinocytes get damaged irreversibly by pollution exposure, they enter senescence which prevents further proliferation. Additionally, the cells are driven into p21^Cip1^-dependent terminal differentiation. This might serve to enhance the skin barrier in the face of external insult. However, when the epidermis is exposed to pollution for extended periods of time, the ongoing accumulation of senescent cells within the basal layer may negatively affect the epidermal and dermal compartment via the SASP (Victorelli et al., 2019, Waldera Lupa et al., 2015, Wang et al., 2017), contributing to the aging process. Pollution-induced skin aging may also occur due to stem cell niche depletion resulting from ongoing DNA damage-induced senescence and late-stage terminal differentiation.

Our study provides novel insights as to how our skin may deal with external aggressors such as air or environmental pollution. However, extensive mechanistic characterisation of pollution-induced senescence and differentiation is needed to better understand how the epidermis reacts to extrinsic stressors. Future work could also focus on targeting the AhR signalling pathway to develop (cosmetic) interventions that ameliorate the consequences of pollution.

## MATERIALS AND METHODS

### Diesel Particulate Matter (DPM), Benzo[a]pyrene (B[a]P) and 2,3,7,8-Tetrachlorodibenzo-p-dioxin (TCDD) preparation

Diesel Particulate Matter SRM 1650b (referred to henceforth as DPM) was chosen as the diesel PM standard to simulate PM_2.5_ exposure. According to National Institute of Standards and Technology (NIST, USA), DPM contained particles with a mean diameter of 0.18µm and was made up mainly of PAHs and nitro-PAHs (Ryu et al., 2019b). In addition, Benzo[*a*]pyrene and 2,3,7,8-Tetrachlorodibenzo-p-dioxin (TCDD) were chosen as Aryl Hydrocarbon Receptor agonists. DPM, B[a]P and TCDD was purchased from Sigma-Aldrich (St. Louis, USA). DPM, B[a]P and TCDD was then dissolved directly in dimethyl sulfoxide (DMSO)at a concentration of 100% and was thoroughly mixed with vortexing and briefly spun down before use. The concentration of DMSO when dissolved in EpiLife™ media and given to cells did not exceed 0.1%.

### Primary keratinocyte culture

Human primary keratinocytes (HPKs) were acquired from the A*STAR Asian Skin Bank (12SKera006) and Cell Research Corp (NK187, NK110). HPKs were cultured and maintained in the presence of mitomycin-C inactivated 3T3/J2 fibroblast feeders in 3:1 [v/v] DMEM (Hyclone, USA)/Ham’s F-12 Nutrient Mix (Gibco, USA) containing 10% Fetal Bovine Serum (Hyclone, USA), 2mM L-glutamine (Sigma-Aldrich, USA), 1nM Cholera Toxin (Sigma-Aldrich, USA), 1µg/ml Hydrocortisone (Sigma-Aldrich, USA), 200ng/ml Recombinant Human EGF (Peprotech, USA) and 10µM Y-27632 dihydrochloride (Tocris Bioscience, UK), otherwise termed as CCMY (Complete culture medium with Y-27632).

3T3-J2 feeder mouse embryonic fibroblast cells were cultured in DMEM supplemented with 10% Fetal Bovine Serum, 1% Penicillin/Streptomycin and 2mM L-glutamine and was inactivated by treatment with 10µg/ml Mitomycin C (STEMCELL Technologies, Canada) for an hour, and allowed to adhere overnight before keratinocytes were seeded onto dishes.

For experimentation purposes, HPKs from CCMY were seeded into EpiLife™ Medium, with 60 µM calcium (Thermo Fisher Scientific, USA) supplemented with Human Keratinocyte Growth Serum (Life Technologies, USA). Keratinocytes were allowed to form colonies before DPM, B[a]P or TCDD treatment and were only used on the third or fourth passage for experimentation. HPKs were treated with 20-30 μg/ml DPM, 2μM B[a]P or 20nM TCDD for 72 hours for experimentation.

All cells were grown at 37°C, 5% CO2 and atmospheric oxygen.

### Senescence-associated beta-galactosidase (SA-**β**-Gal) activity assay

Senescence β-Galactosidase Staining Kit (Cell Signalling Technology, USA) was used according to manufacturer’s protocol and imaged with both brightfield and phase-contrast microscopy. Senescent keratinocytes that turned blue were counted under brightfield microscopy. Total number of keratinocytes was counted using phase-contrast microscopy, and about 300-1000 cells from at least 2-6 different images per condition were counted.

### Real-time Quantitative Reverse Transcription PCR (qRT-PCR)

Total RNA was isolated using RNeasy Mini Kit (Qiagen, Germany) and treated with DNase I as per manufacturer instructions. Isolated RNA was used for reverse transcription using qScript cDNA SuperMix (Quantabio, USA) according to manufacturer’s instructions. qRT-PCR was performed on QuantStudio™ 7 Flex Real-Time PCR System with PowerUp™ SYBR™ Green Master Mix (Applied Biosystems, USA). Analysis of qRT-PCR results were performed in QuantStudio^TM^ Real-Time PCR Software (Applied Biosystems, USA). Amplification values were normalised to beta-actin, and primers used is shown in Table 1.

**Table 1.**
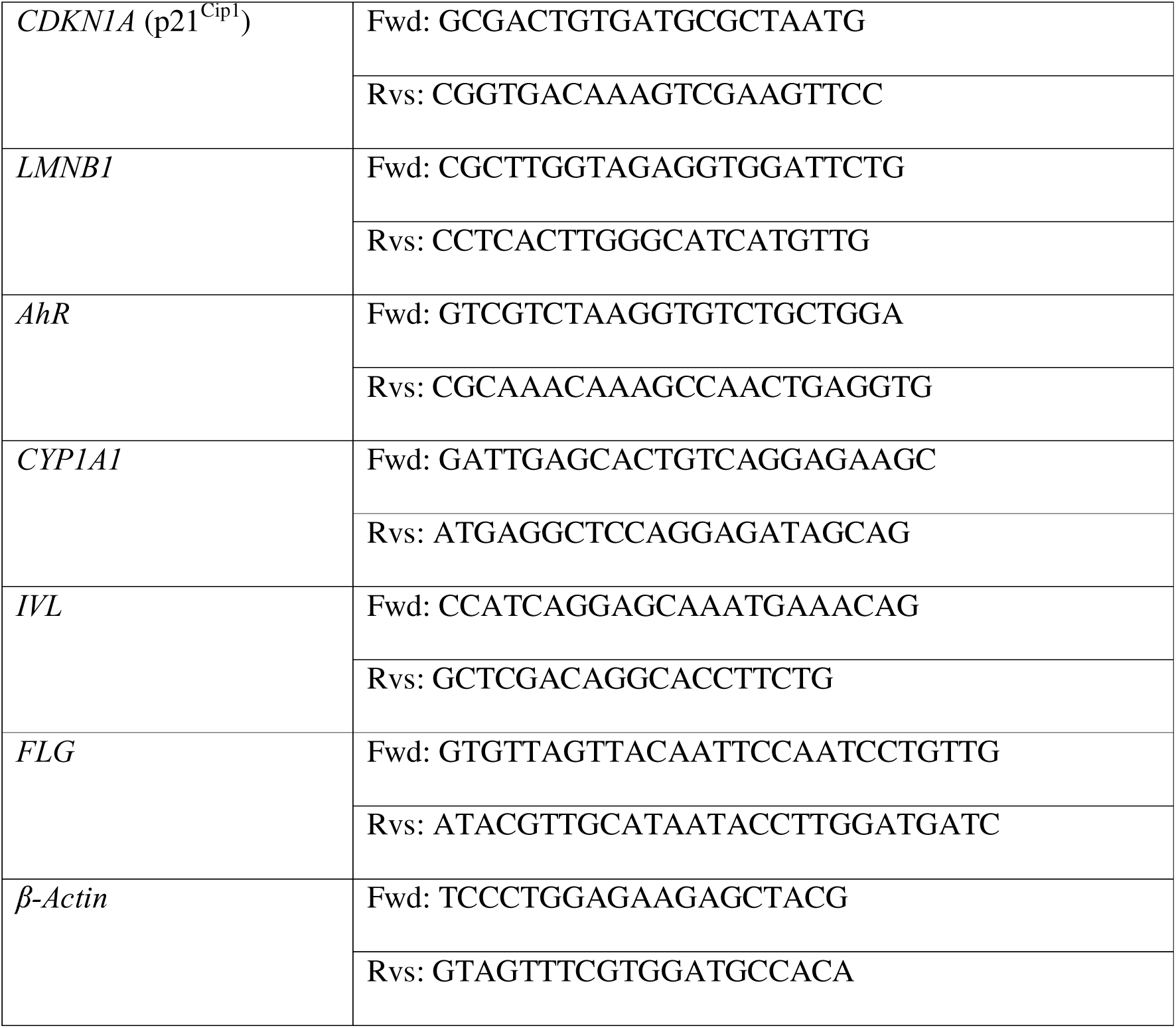
Primers used for qRT-PCR.

### Transcriptomics and bioinformatics analysis

Total RNA was isolated from human primary keratinocytes using RNeasy Mini Kit (QIAGEN, Germany) and treated with DNase I as per manufacturer instructions. All experiments were done in triplicate (DMSO CTRL vs 2μM B[a]P), making up a total of 6 samples. The RNA integrity was assessed with an Agilent 2100 Bioanalyzer (Agilent Technologies).

Differential expression analysis of two conditions/groups (two biological replicates per condition) was performed using the DESeq2Rpackage (1.20.0). The resulting P-values were adjusted using the Benjamini & Hochberg approach for controlling the false discovery rate (FDR). Genes with an adjusted P-value <=0.05 found by DESeq2 were assigned as differentially expressed. Prior to differential gene expression analysis, for each sequenced library, the read counts were adjusted by EdgeR program package through one scaling normalized factor. Differential expression analysis of two conditions was performed using the edgeR R package (3.22.5). The p values were adjusted using the Benjamini & Hochberg method. Corrected p-value of 0.05 and absolute foldchange of 2 were set as the threshold for significantly differential expression.

Functional enrichment analysis was performed using Gene Ontology (GO) and Gene Set Enrichment Analysis (GSEA). GO enrichment analysis of differentially expressed genes was implemented by the clusterProfiler R package, in which gene length bias was corrected. GO terms with corrected p-value less than 0.05 were considered significantly enriched by differential expressed genes. GSEA was utilised to rank gene sets according to the degree of differential expression in the two samples, and then the predefined Gene sets were tested to see if they were enriched at the top or bottom of the list. FDR of <=0.25 was taken as a significant gene set enrichment.

### Immunofluorescence

HPKs were seeded onto μ-Slide 8 Well IbiTreat chamber slides (Ibidi Germany) and allowed to form mini-colonies. HPKs were fixed at 2% paraformaldehyde for 15 minutes, and permeabilised with 0.1% Triton X-100 in 1X PBS for 5 minutes. The cells were then blocked with 10% Donkey Serum (Sigma-Aldrich, USA) in 1X PBSA (0.5% BSA, 0.1% Saponin, 0.01% Sodium Azide dissolved in 1X PBS) for 60 minutes and were incubated with primary antibodies diluted in 1X PBSA with 2% Donkey Serum overnight. Following primary antibody incubation, the HPKs were incubated in secondary antibodies diluted in 1X PBSA with 2% Donkey Serum for 60 minutes. Cells were stained and mounted in 1µg/ml DAPI diluted in Fluoromount Aqueous Mounting Medium (Sigma-Aldrich, USA) and imaged using the FV3000RS Olympus Confocal Laser Scanning Microscope (x20 and x60 magnification for representative images, x20 magnification for quantification) and analysed with a custom macro on FIJI, ImageJ. Primary and secondary antibodies are listed in tables 2 and 3, respectively.

**Table 2.**
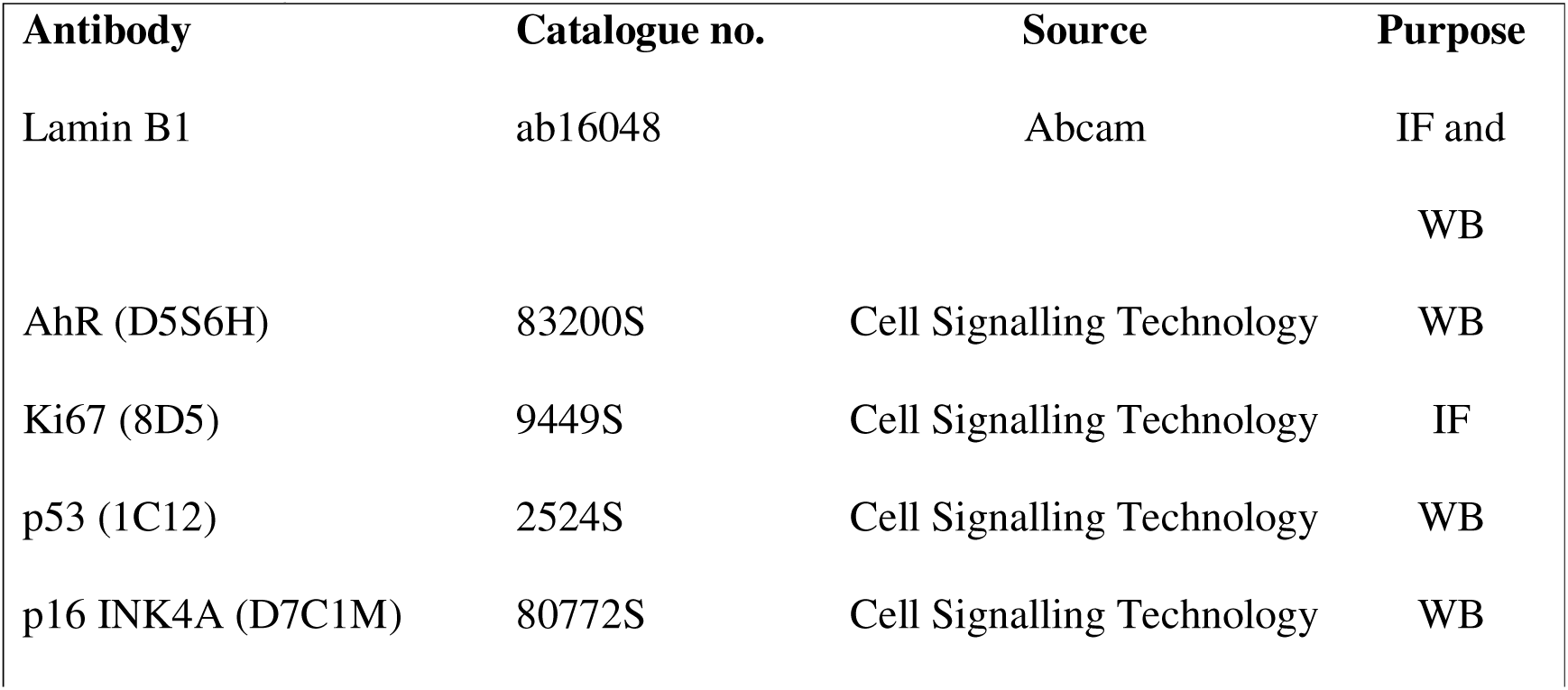
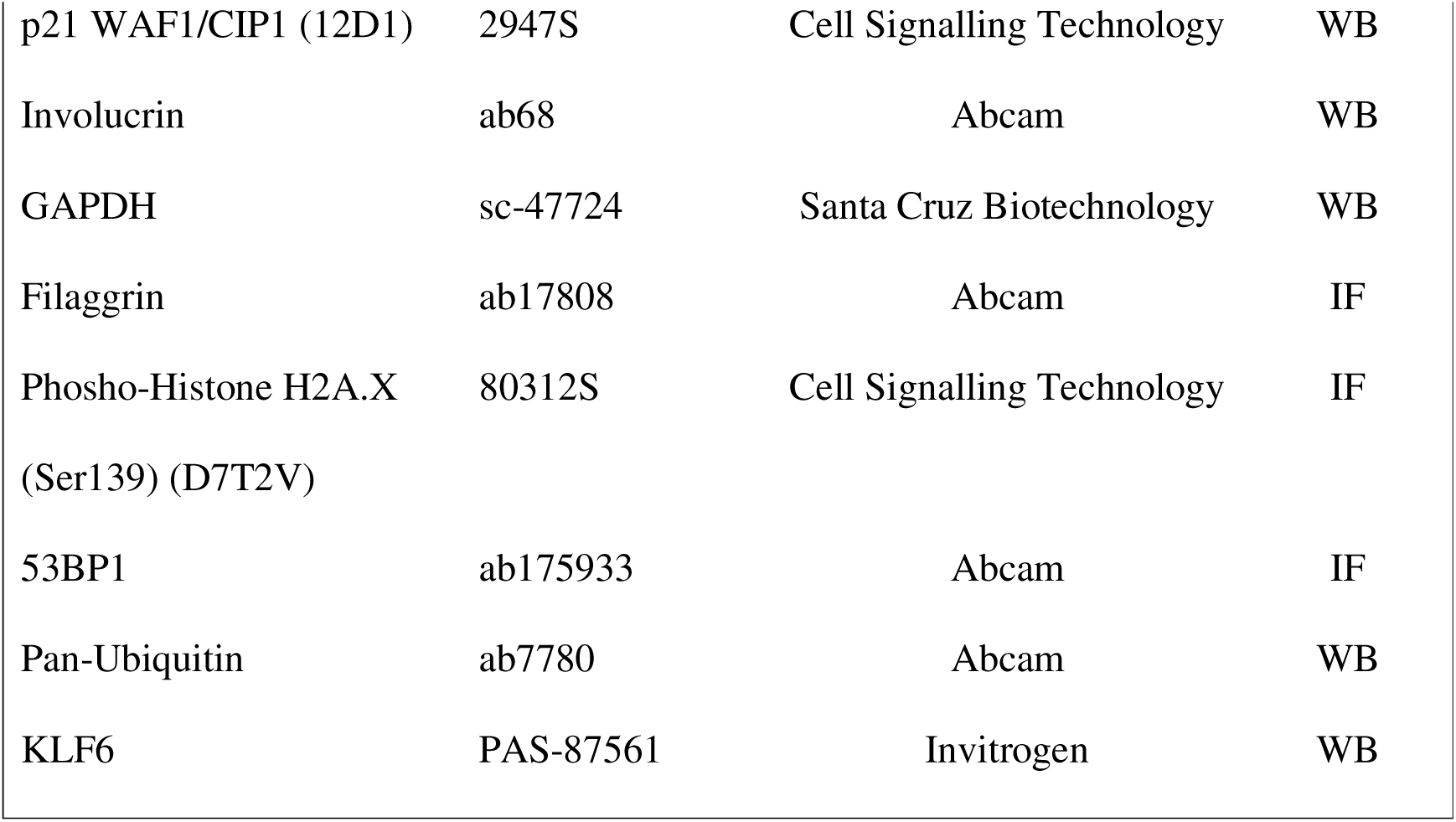
Primary antibodies used.

| **Antibody** | **Catalogue no.** | **Source** | **Purpose** |
| --- | --- | --- | --- |
| Lamin B1 | ab16048 | Abcam | IF and<br>WB |
| AhR (D5S6H) | 83200S | Cell Signalling Technology | WB |
| Ki67 (8D5) | 9449S | Cell Signalling Technology | IF |
| p53 (1C12) | 2524S | Cell Signalling Technology | WB |
| p16 INK4A (D7C1M) | 80772S | Cell Signalling Technology | WB |
| p21 WAF1/CIP1 (12D1) | 2947S | Cell Signalling Technology | WB |
| Involucrin | ab68 | Abcam | WB |
| GAPDH | sc-47724 | Santa Cruz Biotechnology | WB |
| Filaggrin | ab17808 | Abcam | IF |
| Phospho-Histone H2A.X<br>(Ser139) (D7T2V) | 80312S | Cell Signalling Technology | IF |
| 53BP1 | ab175933 | Abcam | IF |
| Pan-Ubiquitin | ab7780 | Abcam | WB |
| KLF6 | PAS-87561 | Invitrogen | WB |

**Table 3.** Secondary antibodies used.

| **Antibody** | **Catalogue no.** | **Source** | **Purpose** |
| --- | --- | --- | --- |
| Donkey Anti-Rabbit IgG<br>H&L Alexa Fluor 568 | Ab175470 | Abcam | IF |
| Donkey Anti-Mouse IgG<br>H&L Alexa Fluor 488 | Ab150105 | Abcam | IF |
| IRDye 800CW Donkey<br>anti-Mouse IgG<br>Secondary Antibody | 926-32212 | LI-COR Biosciences | WB |
| IRDye 680RD Goat anti-Rabbit IgG Secondary<br>Antibody | 926-68071 | LI-COR Biosciences | WB |

### Western Blotting

Whole cell lysis was performed using a lysis buffer consisting of 25mM HEPES, 25mM MgCl_2_, 300mM NaCl, 0.1mM EDTA, 20% Glycerol and 0.2% Sodium Dodecyl Sulfate, and sonicated at low frequency for 20 cycles at 30 seconds on/off cycles.

Protein was quantified with DC Protein Assay (Bio-Rad Laboratories, USA). Equal amounts of protein were separated by SDS-PAGE before transfer and western blotting. Fluorescent protein bands were visualised using the LI-COR’s Odyssey Infrared Imaging System according to manufacturer’s protocol.

### Generation of shRNA knockdown lentivirus and lentiviral transduction

shRNAs targeted against AhR (shAhR 1A and 2A), p21 (shp21) and KLF6 (shKLF6) were cloned into the pLKO.1-puro plasmid (Addgene, #8453) via restriction cloning. The following plasmids were transfected into HEK293T cell line with pCMV-VSV-G (Addgene #8454), pCMV delta R8.2 (Addgene, #12263) and the respective pLKO.1 plasmids at 2.5μg: 2.5μg:10μg respectively. Lentivirus was collected daily for four days and concentrated using Lentiviral Concentrator (OriGene, TR30025). Lentiviral transduction was performed in EpiLife media, with 5μg/ml polybrene over the span of 24 hours. Puromycin selection (5μg/ml) was carried out on the fourth day for 24 hours and were split afterwards for either experiments or maintenance in CCMY.

### Histology

Tissues were fixed in 4% paraformaldehyde for 4 hours at room temperature, followed by overnight processing of tissues using the Leica Automatic Tissue Processor (Leica Biosystems, Germany), according to manufacturer’s protocol. Tissues were then embedded in paraffin the next day, sectioned and mounted onto polysine microscope slides (Thermo Fisher Scientific, USA). Hematoxylin and Eosin (H&E) was performed using the Leica AutoStainer (Leica Biosystems, Germany), and imaged using the Zeiss AxioScan.Z1 Slide Scanner (Zeiss, Germany) with a 20x objective lens. Prior to immunofluorescence staining on sections, antigens were retrieved by baking slides for one hour at 42[C, followed by clearene incubation and ethanol washes, and then placing the tissue sections in pH 6.0 Sodium Citrate Buffer for 20 minutes at 90[C. Sections were then transferred to PBS with sodium azide.

### Statistical testing

P-values for all non-transcriptomic analysis were obtained using ANOVA one-way analysis through GraphPad Prism 8. Results are displayed as mean ±[SD in figures unless stated otherwise. p<0.05 values were considered significant. (*p[<[0.05, **p[<[0.01, ***p[<[0.001, ****p<0.0001).

### Ethics statement

N/A

## Data availability statement

RNAseq data have been deposited in the GEO database (https://www.ncbi.nlm.nih.gov/geo/) under accession number xxx. All other data are available upon request.

## Acknowledgments

This work was funded by a Singapore Economic Development Board- Industrial Postgraduate Programme (EDB-IPP) grant, jointly funded with Kenvue, Inc. The funders did not have any involvement in the work.

## CrediT statement

Conceptualization: MvS, MT; Methodology: DL, MvS; Investigation: DL; Data Curation: DL; Writing - Original Draft: DL; Writing - Review & Editing: MvS, DL; Visualization: DL; Supervision: MT, MvS; Funding acquisition: MvS, MT

## Declaration of AI/LLM

No AI/LLM tools were used to prepare this manuscript.

## SUPPLEMENTARY FIGURE LEGENDS

**Supplementary Figure S1:**
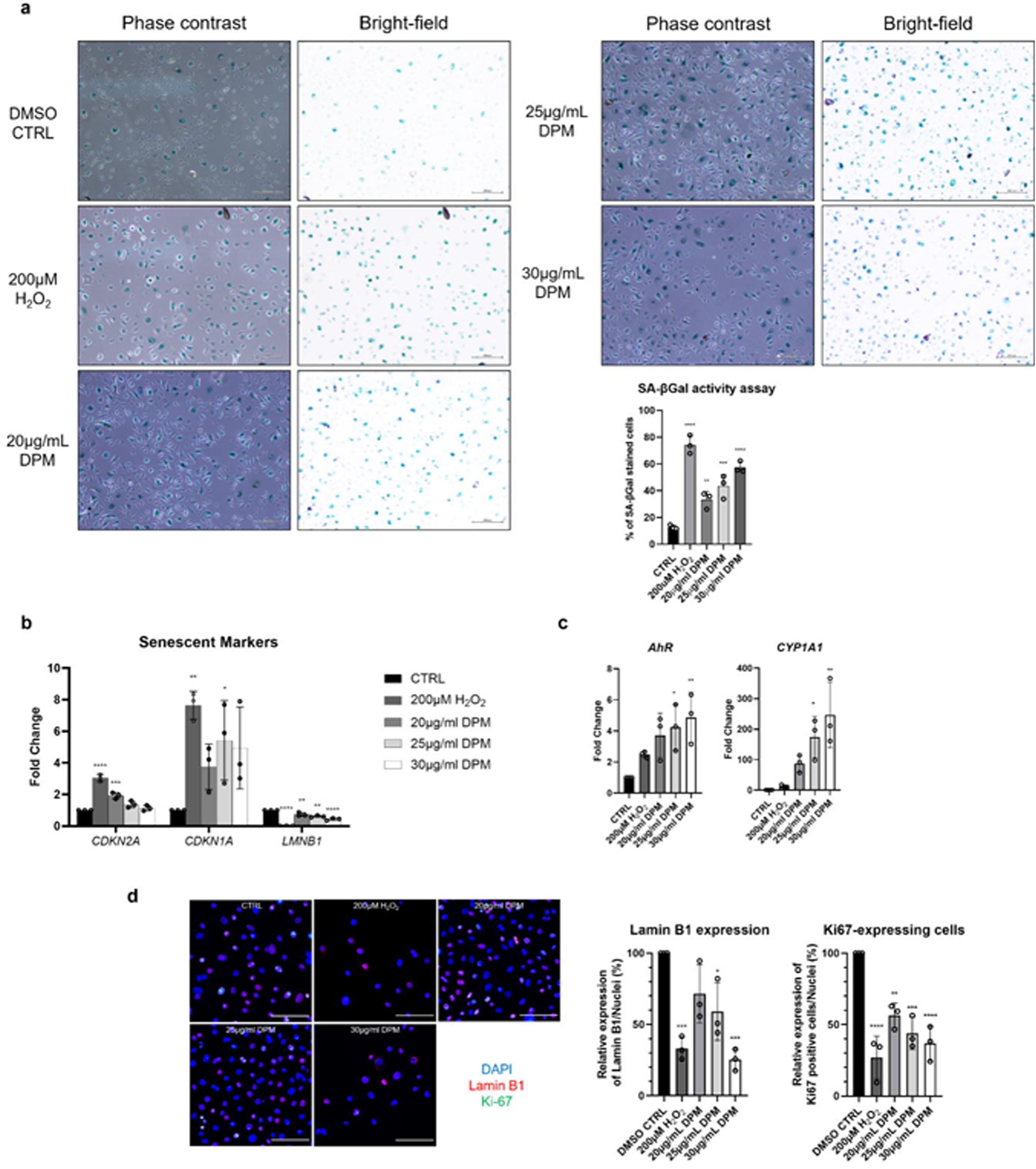
Diesel Particulate Matter induces cellular senescence in primary keratinocytes. (a) SA-β-Gal staining of HPKs (2D cultures) treated with HPKs with 200μM H_2_O_2_ for 1h, then allowed to recover for 4 days or with 20-30μg/ml DPM for 4 days. Bar = 200μm. (b) RT-qPCR analyses for senescence-associated genes after treating HPKs as in a. (c) RT-qPCR of AhR effector genes after treating HPKs as in a. (d) Immunofluorescence analyses (Lamin B1, Red; Ki67, Green; DAPI, Blue) after treating HPKs as in a. Bar = 100μm. All treatments were compared to the treatment control for statistical testing. Statistical analyses: One-way ANOVA, n = 3, *P*-values < 0.05 indicate statistical significance (∗P < 0.05; ∗∗P < 0.01; ∗∗∗P < 0.001; ∗∗∗∗P < 0.0001). SA-β-Gal (Senescence-Associated Beta Galactosidase); H_2_O_2_ (Hydrogen peroxide); 2D (two-dimensional); DPM (Diesel Particulate Matter); HPK (Human Primary Keratinocyte); *LMNB1* (Lamin B1); *CDKN1A* (p21^Cip1^).

**Supplementary Figure S2:**
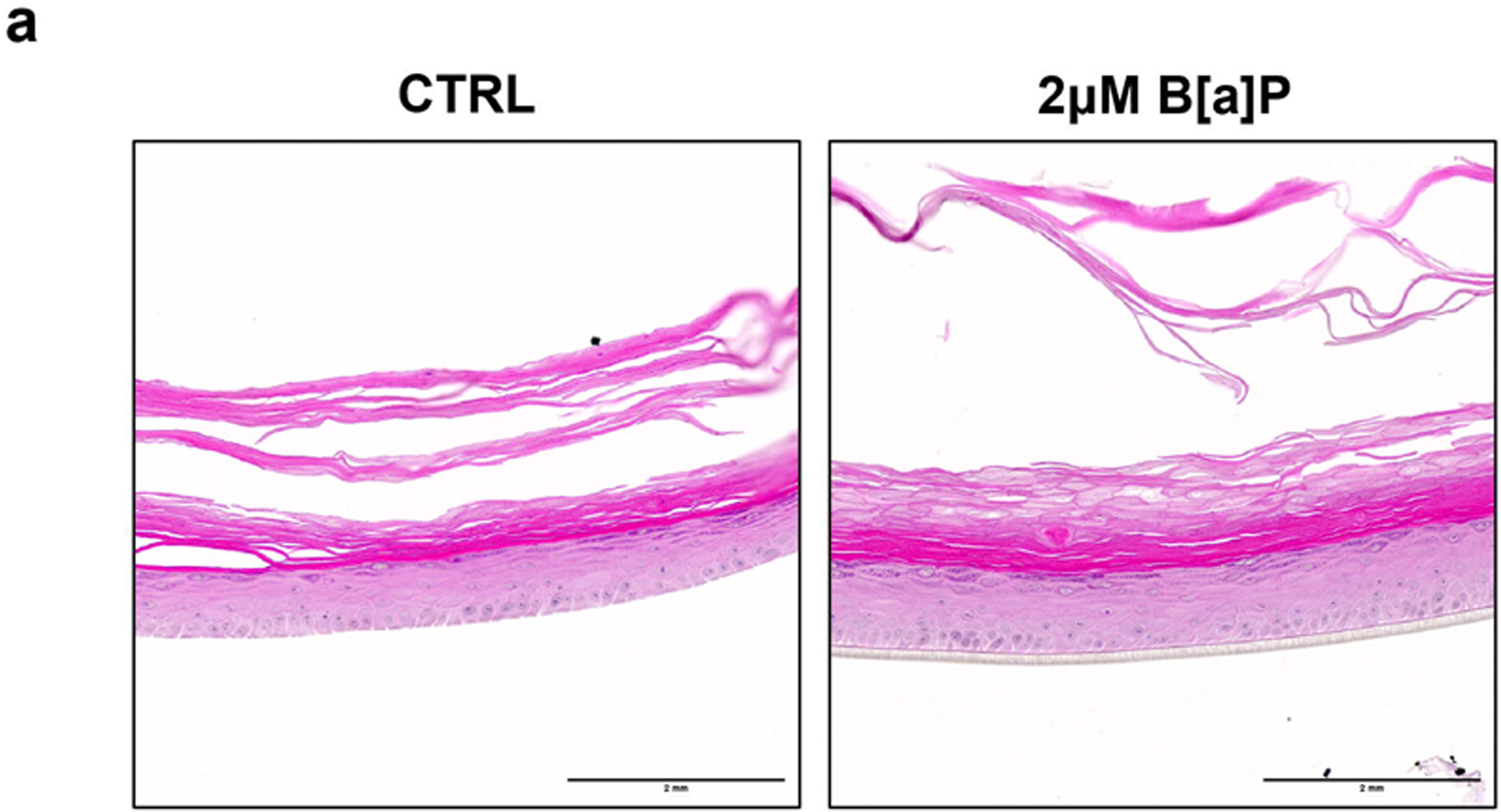
Benzo[a]pyrene does not induce epidermal morphological changes in RHEs. (a) Hematoxylin and Eosin (H&E) staining of 3D Reconstructed Human Epidermis organotypic cultures (RHEs) after 7 days of 2μM B[a]P exposure or 0.1% DMSO (CTRL). Scale bar indicates 2mm.

**Supplementary Figure S3:**
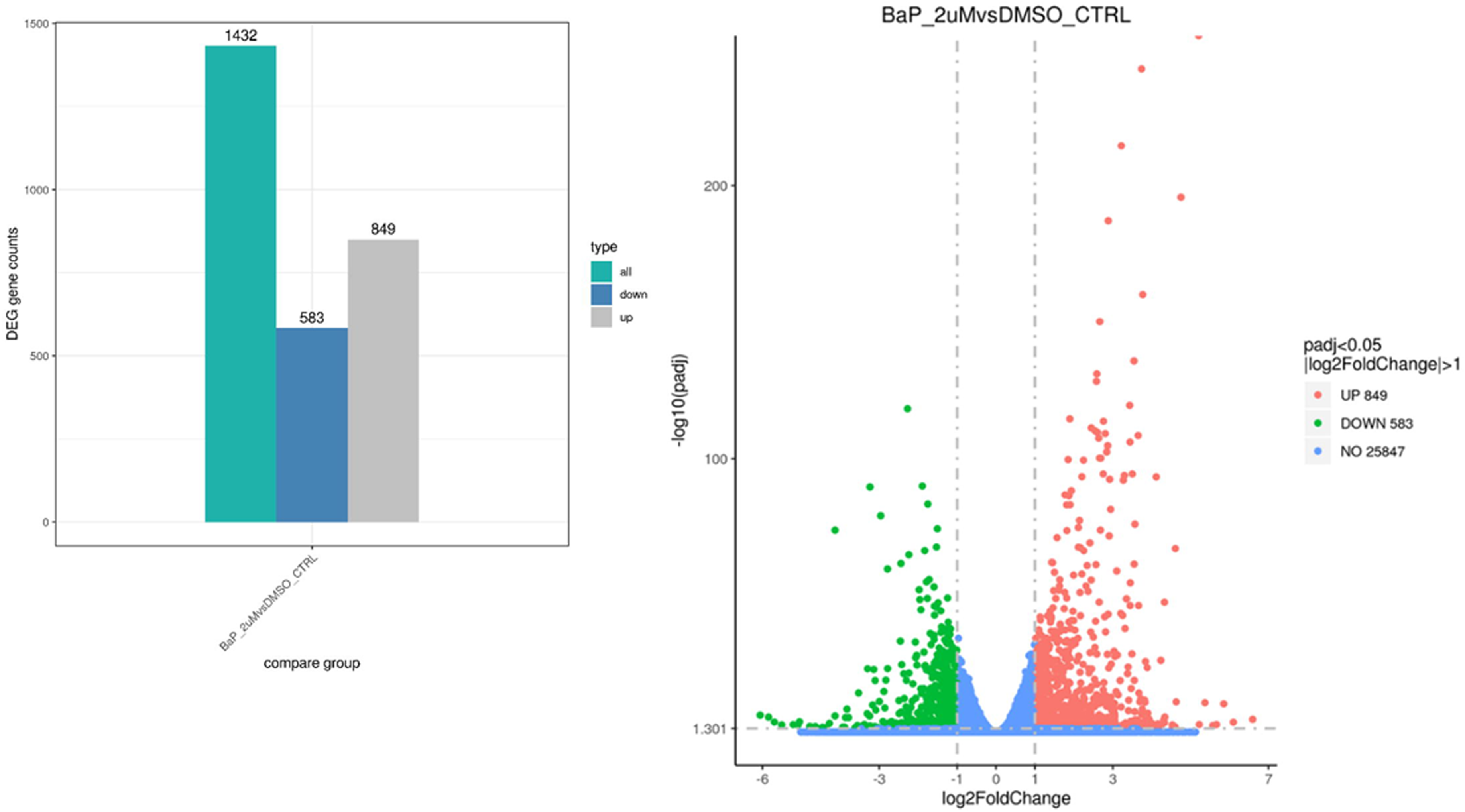
Significantly differentially expressed genes in 2μM B[a]P-exposed human primary keratinocytes (HPK). (a) Significantly differentially expressed genes between 2μM B[a]P (96 hours) versus control (0.1% DMSO). (b) Volcano plot depicting the log_2_Foldchange against -log_10_(adjusted *p-*value), significantly differentially expressed genes were defined as adjusted *p*-value<0.05 and |log_2_FoldChange|>1.

**Supplementary Figure S4:**
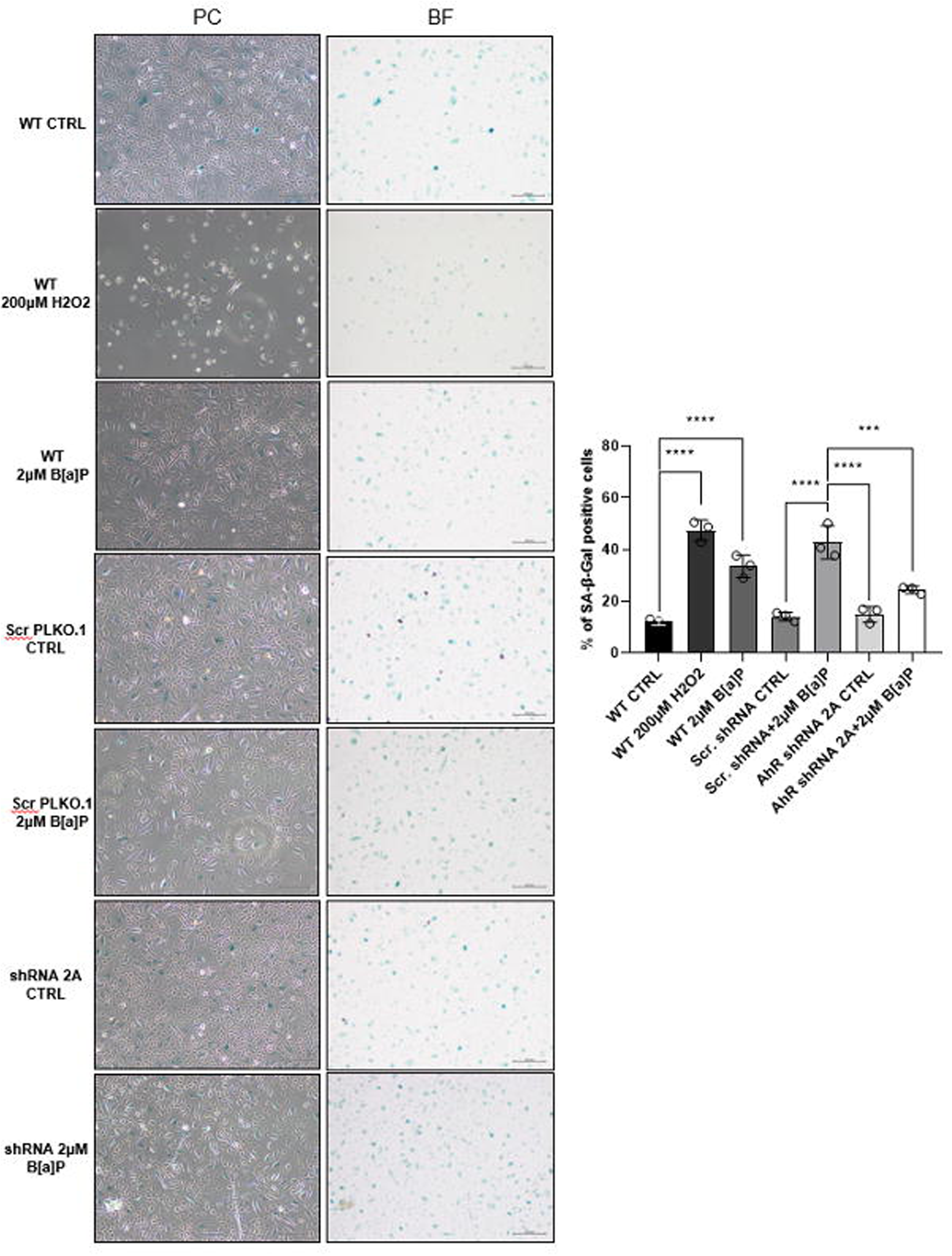
Partial AhR knockdown rescues HPKs from Benzo[a]pyrene-induced senescence. SA-β-Gal assay performed after treating wild-type HPKs with 200μM H_2_O_2_ for 1 hour and allowed 4 days to recover and 2μM B[a]P for 4 days, as well as Scrambled PLKO.1- and shRNA 2A-transduced HPKs treated with 2μM B[a]P for 4 days. Bar = 200μm. Statistical analyses: One-way ANOVA, n = 3, *P*-values < 0.05 indicate statistical significance (∗P < 0.05; ∗∗P < 0.01; ∗∗∗P < 0.001; ∗∗∗∗P < 0.0001).

**Supplementary Figure S5:**
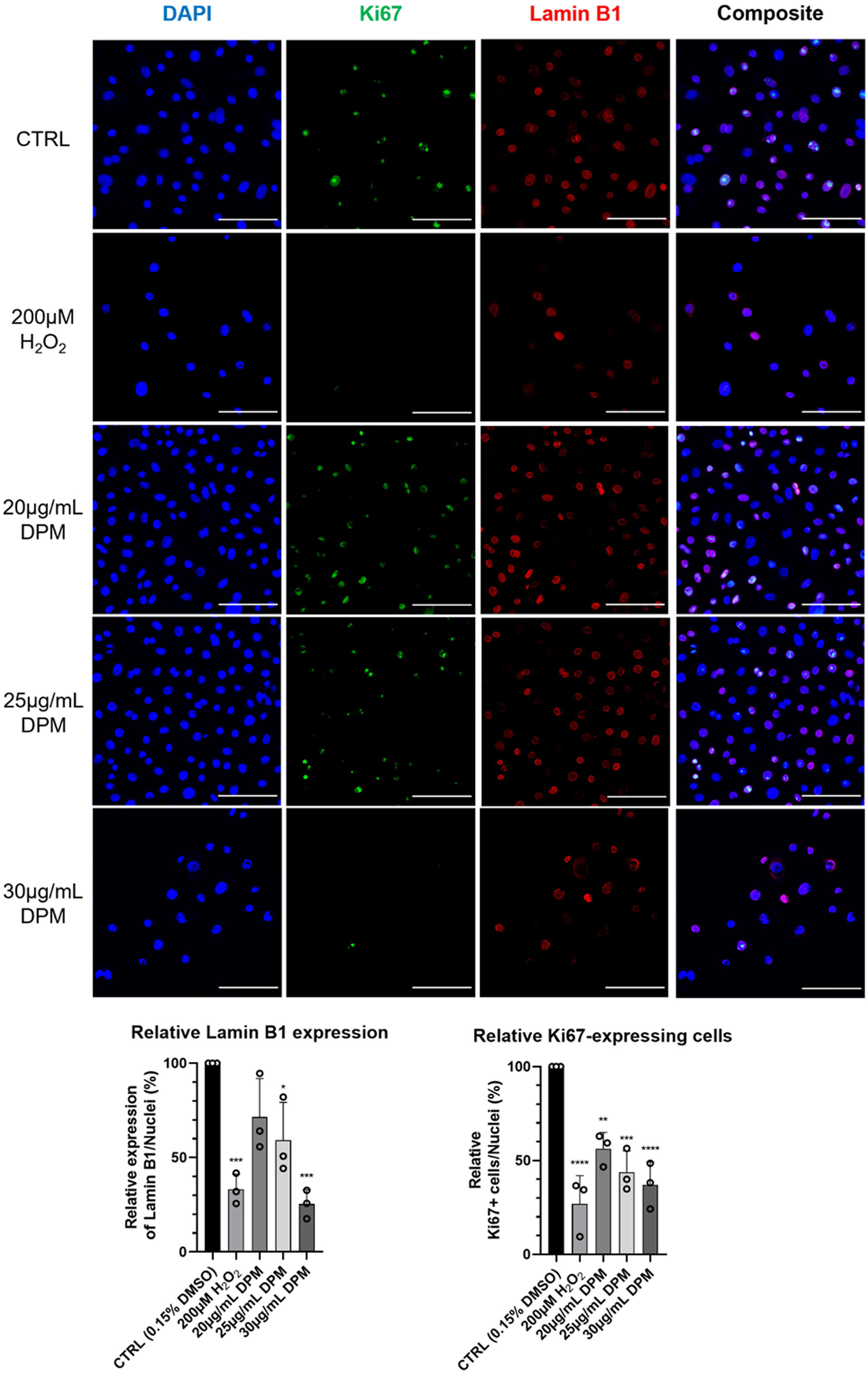
Diesel Particulate Matter induces cellular senescence in primary keratinocytes. Complete panels of immunofluorescence analyses from Supplementary Figure S1d (Lamin B1, Red; Ki67, Green; DAPI, Blue) of HPKs (2D cultures) treated with 200μM H_2_O_2_ for 1h, then allowed to recover for 4 days or with 20-30μg/ml DPM for 4 days. Bar = 100μm. All treatments were compared to the treatment control for statistical testing. Mean fluorescence intensity was plotted for 3 biological replicates. Statistical analyses: One-way ANOVA, n = 3, *P*-values < 0.05 indicate statistical significance (∗P < 0.05; ∗∗P < 0.01; ∗∗∗P < 0.001; ∗∗∗∗P < 0.0001).

**Supplementary Figure S6:**
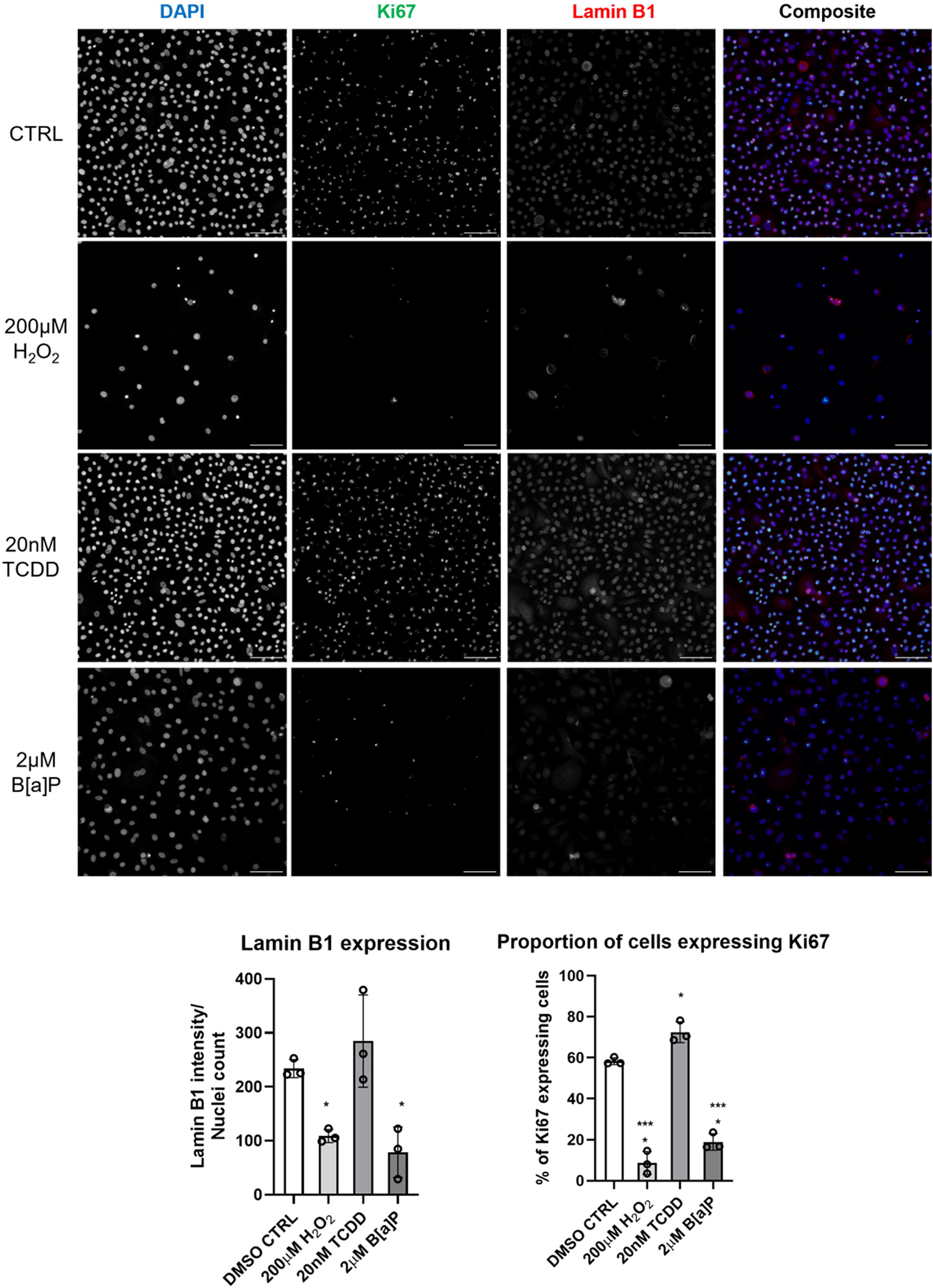
Benzo[a]pyrene induces cellular senescence in primary keratinocytes. Complete panels of immunofluorescence analyses from Figure 1d (Lamin B1, Red; Ki67, Green; DAPI, Blue) of HPKs (2D cultures) treated with HPKs with 200μM H_2_O_2_ for 1h and allowing 5 days for recovery, and 20nM TCDD and 2μM B[a]P for 5 days. Bar = 100μM. All treatments were compared to the treatment control for statistical testing. Mean fluorescence intensity was plotted for 3 biological replicates. Statistical analyses: One-way ANOVA, n = 3, *P*-values < 0.05 indicate statistical significance (∗P < 0.05; ∗∗P < 0.01; ∗∗∗P < 0.001; ∗∗∗∗P < 0.0001).

**Supplementary Figure S7:**
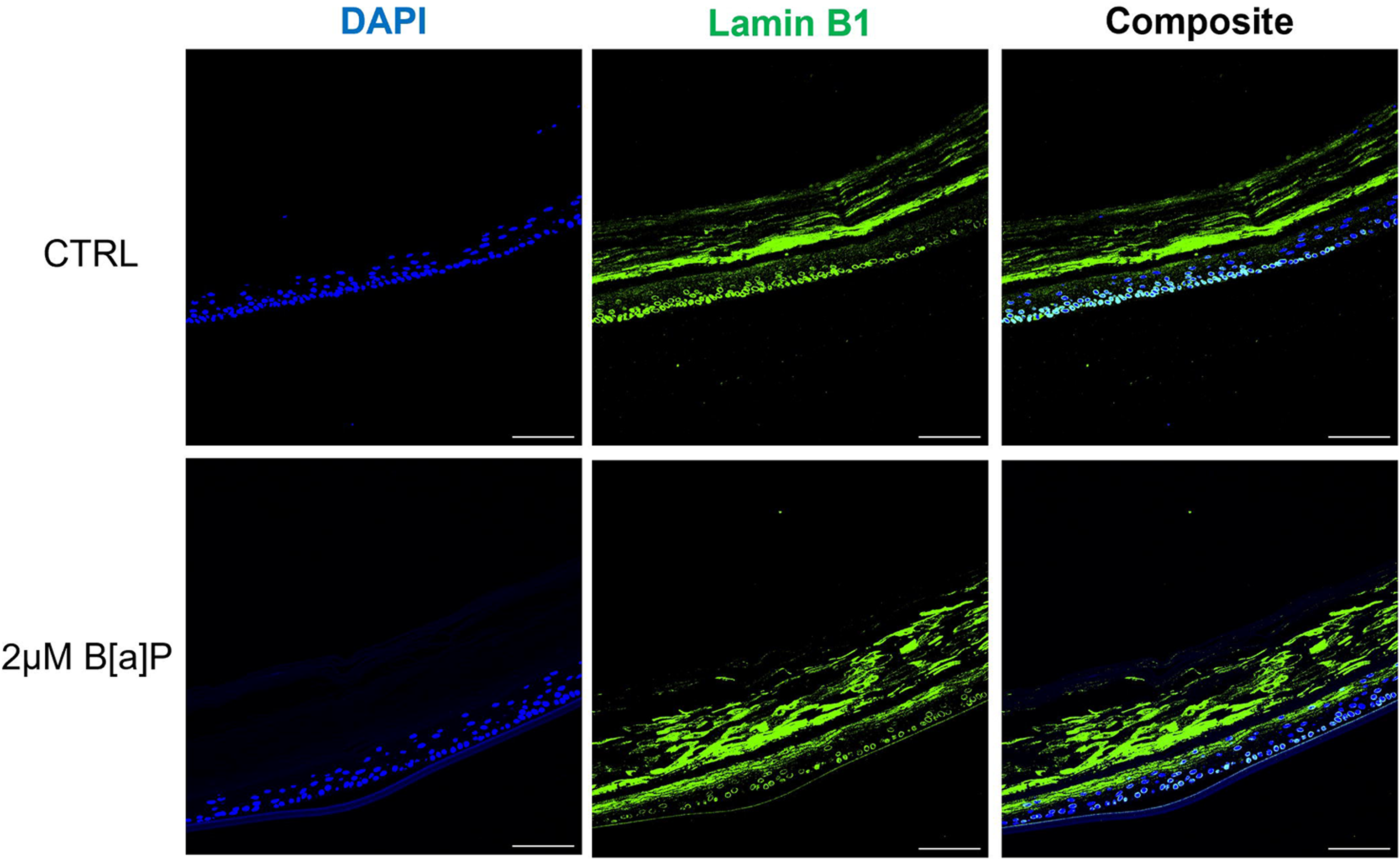
Benzo[a]pyrene induces cellular senescence in reconstructed human epidermis. Complete panels of immunofluorescence analyses from Figure 1e (Lamin B1, Green; DAPI, Blue) of Reconstructed Human Epidermis topically treated with 2μM B[a]P or DMSO (CTRL) after 7 days. Scale bar = 100μm.

**Supplementary Figure S8:**
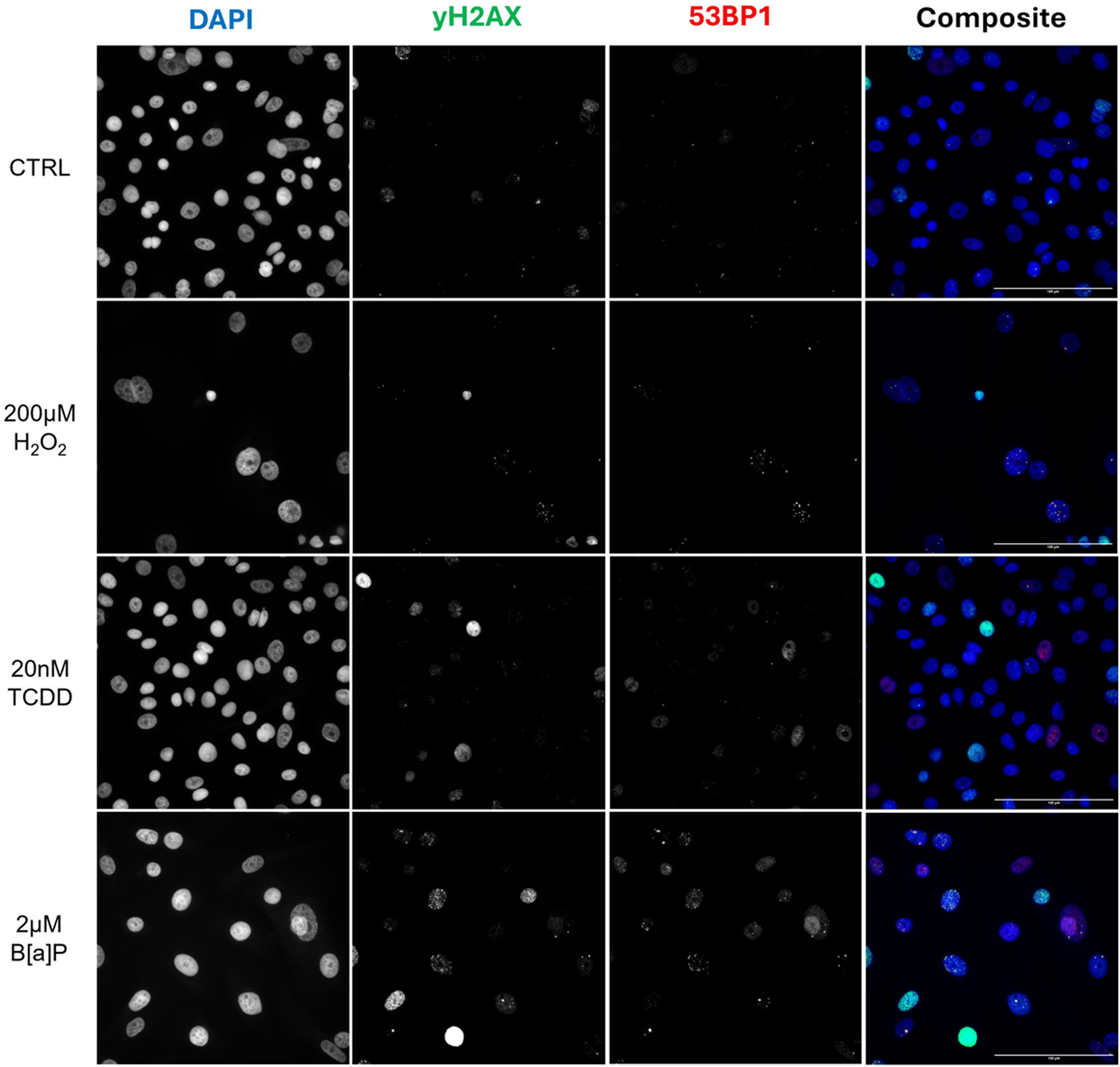
Benzo[a]pyrene induced DNA damage in primary keratinocytes. Complete panels of immunofluorescence analyses from Figure 1f (53BP1, Red; γH2AX, Green; DAPI, Blue) was performed after treating HPKs with 200μM H2O2 for 1h and allowed 5 days for recovery, and 20nM TCDD and 2μM B[a]P for 5 days. White arrows indicate DNA damage foci. Bar = 100μm. All treatments were compared to DMSO control for statistical testing. Statistical analyses: One-way ANOVA, n = 3, P-values < 0.05 indicate statistical significance (∗P < 0.05; ∗∗P < 0.01; ∗∗∗P < 0.001; ∗∗∗∗P < 0.0001).

**Supplementary Figure S9:**
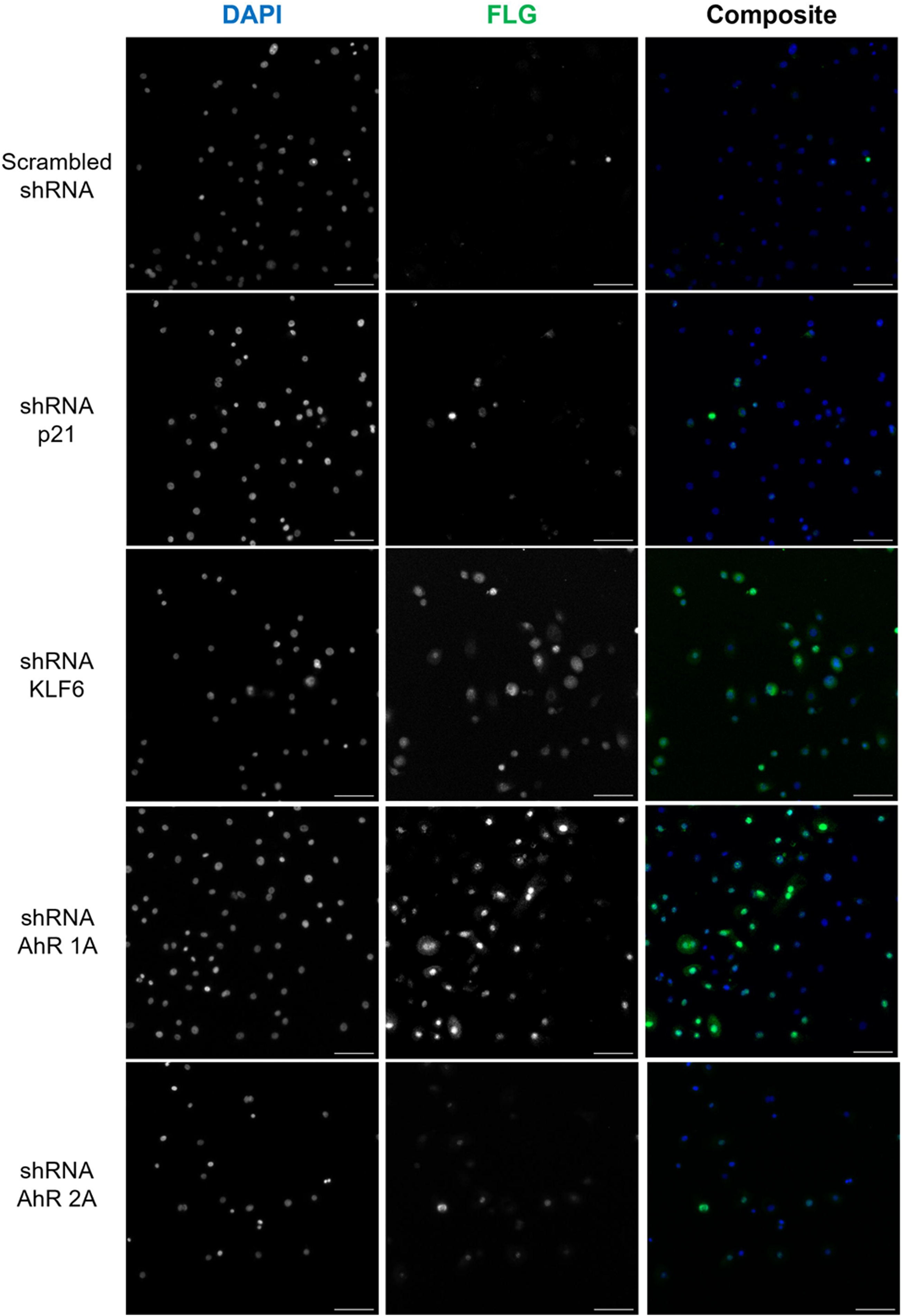
Complete knockdown of AhR and partial knockdown of p21Cip1 induced filaggrin expression in primary keratinocytes. Complete panels of immunofluorescence analyses of near complete shRNA-mediated knockdown of AhR (shAhR 1A), Partial shRNA-mediated knockdown of AhR (shAhR 2A), Partial shRNA-mediated knockdown of p21^Cip1^ (shp21), Partial shRNA-mediated knockdown of KLF6 (pKLF6) was carried out 4 days post-lentiviral transduction from Figure 4c (Filaggrin, Green; DAPI, Blue) was carried out. Bar = 100μm

